# Genomic and environmental determinants of circular single-stranded DNA production in engineered *E. coli*

**DOI:** 10.64898/2026.10.28.761454

**Authors:** Konlin Shen, Shawn M. Douglas

**Affiliations:** Department of Cellular and Molecular Pharmacology, University of California, San Francisco, San Francisco, CA 94158, USA

## Abstract

Single-stranded DNA with hundreds to thousands of bases is a critical reagent in biotechnology. While traditional chemical synthesis is not suitable for efficient production of such long oligonucleotides, M13 bacteriophage offers a scalable pathway towards custom, long, single-stranded DNA production. However, conventional helper plasmid-phagemid systems suffer from batch-to-batch variability and off-target packaging. To address these limitations, we previously engineered eScaf, an E. coli strain with the M13 genome stably integrated into its chromosome. eScaf produces circular, single-stranded DNA with high purity, but yields of only 100 and 200 µg of DNA per liter of culture, too low for cost-effective scale-up. Here, we systematically investigate the genomic and environmental determinants of cssDNA yield in eScaf, revealing that cssDNA yield depends on both the number and chromosomal loci of integrated M13 genomes. Combining these insights with optimized media, we achieve ∼4 mg/L of cssDNA in shake flasks, a 40-fold improvement. We also evaluate fed-batch bioreactor feeding strategies for cssDNA production using eScaf. Our new strain, eScaf-IS3, enables cost-effective production of cssDNA for high-throughput therapeutic screening, operon-length knock-in donor production, and large-scale DNA origami assembly.

## Introduction

Synthesizing arbitrary length and sequence DNA oligonucleotides is a non-trivial challenge. Single-stranded DNA (ssDNA) underlies nearly every methodological advance in modern biotechnology, from gene synthesis and genomic knock-ins to structural DNA nanotechnology and nucleic acid therapeutics. While short oligonucleotides are routinely produced by chemical synthesis, this approach breaks down at kilobase lengths due to cumulative errors at each base incorporation step. Many labs are exploring alternative approaches to oligo synthesis (1, 2, 3), notably through enzymatic extension as opposed to direct chemical incorporation, but thus far they are still limited in efficiency and scale. Improving scalability of arbitrary length and sequence oligonucleotides would greatly increase accessibility of nucleic acid products by reducing the cost of long and complex ssDNA strands, leading to innovation in therapies and other applications.

Microbial production of ssDNA using M13mp18 is scalable, but customizability comes at the cost of purity and yield. Rather than using enzymatic or chemical production, harnessing the power of microbes to produce DNA at scale is a tried-and-true solution. To produce ssDNA bacteriologically, M13 bacteriophage, which infects *E. coli*, is grown in bacteria and harvested for its circular ssDNA (cssDNA) genome. The cssDNA genome is replicated inside *E. coli* by the endogenous DNA replication machinery and packaged by the foreign M13 proteins expressed by the *E. coli* after infection. To alter the sequence of the packaged cssDNA, a phagemid encoding the target sequence flanked by the M13 packaging sequence is transformed into *E. coli* and infected with M13 helper phage or co-transformed with M13 helper plasmid to produce M13 particles with the target sequence enclosed (4). Both these methods introduce risk of off-target packaging, either from the helper phage genome or variability in helper plasmid expression and come at the cost of reduced cssDNA yield (5).

To address these purity limitations, we previously constructed an engineered *E. coli* strain, eScaf, by integrating a single copy of the M13mp18 genome at the ulaD locus in *E. coli* TG1 (6). eScaf produces high-purity cssDNA with lengths over 20,000 ntand has been validated for use in high-efficiency genomic editing of primary human T cells. However, yields from eScaf averaged between 100 and 200 µg/L, with a maximum of 250 µg/L, substantially below what helper plasmid-phagemid systems achieve in bioreactors (2-3 mg/L; 7). These yields are also far below those reported for optimized M13 genome production (up to 600 mg/L) (8), or for custom cssDNA sequence production using custom-M13 particle infection (100-200 mg/L) (9)). For important therapeutic applications such as DNA origami vaccine platforms, a single dose may require 50 µg of cssDNA scaffold (10). In T-cell engineering for CAR T-cell therapy, one of the most significant advances in therapeutic technology in the past two decades, 1∼5 µg of cssDNA donor is typically used per 1 M cells (11, 12); taking knock-in efficiency and expansion rate into account, a single patient might need tens of micrograms of cssDNA donor to get ∼50M to 100M cells for infusion. Thus, bringing the yield of eScaf to the milligram-per-liter scale and above would facilitate access to critical cssDNA reagents used for life-saving clinical-scale therapeutics.

Here we systematically investigate the genomic and environmental determinants of cssDNA yield in eScaf, examining the effects of chromosomal copy number, integration locus, host strain genotype, coat protein modification, media composition, and bioreactor feeding strategy. Together these investigations increased cssDNA yield by nearly 40x while retaining the purity and consistency advantages of chromosomal integration.

## Materials and Methods

### Producing cssDNA in shake flasks using bacteria

Unless otherwise specified, strain construction, flask culture conditions, and cssDNA purification and measurement followed our previously described methods (6). To produce cssDNA using bacteria in flasks, we transformed a phagemid containing the desired sequence into eScaf. For testing cssDNA production using His-tagged M13mp18 donors, the M13mp18 donor plasmid was co-transformed with the phagemid as a source of M13 proteins. Transformants were picked and grown in 1 mL LB preculture with antibiotics at 30 °C for at least 8 hours before inoculating the primary culture (target media + antibiotic selection) with preculture. The volume of preculture added to the primary culture was 1/1000^th^ of the primary culture volume (i.e. 50 µL of preculture for a 50 mL flask). The culture was allowed to grow for at least 12 hours at 30 °C before phage particles were harvested and the cssDNA extracted.

### Measuring growth of bacteria

We measured growth by reading the OD600 of bacterial cultures (Ultrospec10). If the OD600 for the culture exceeded 1, we remeasured across three different dilutions for which OD600 of the culture was between 0.1 and 1. A linear fit to the OD600 vs dilution factor data was performed and the OD600 for the original culture at 1x dilution was extrapolated.

### Harvesting and purifying cssDNA products from bacterial culture

Bacterial cultures ready for harvest were chilled on ice for 35’, then pelleted at 7000 RCF, 4 °C for 15’. The supernatant containing the M13 particles was decanted into clean tubes and bacterial pellets discarded. The supernatant was then mixed with 0.25 volumes of a 5x PEG fractionation buffer (200 g/L PEG8000, 150 g/L NaCl) and the phage were left to precipitate on ice for 30’. The phage particles were pelleted by centrifugation at 9000 RCF, 4 °C for 15’. The supernatant was carefully discarded, and the phage particles were resuspended in TE buffer.

The resuspended phage were spun for 15’ at 15 kRCF, 4 °C to pellet any residual bacteria. The supernatant containing the phage was moved to a clean tube, then spun again for 15’ at 15 kRCF, 4° C for a second clarifying spin. The supernatant was once again moved to a clean tube and the tube containing the pellet discarded.

Finally, to extract the cssDNA from the phage particles, the 2 volumes of lysis buffer (P2, Qiagen) were added to the phage solution followed by 1.5 volumes of neutralization buffer (P3, Qiagen). The mixture was incubated on ice for 15’, then centrifuged for 15’ at 15 kRCF, 4 °C. The supernatant containing the DNA was moved to a clean tube and 2 volumes of 100% ethanol were added to the mixture to precipitate the DNA. The DNA-ethanol mixtures were allowed to sit at room temperature for at least 30’, then the DNA was pelleted for 30’ at 16000 RCF at room temperature. The supernatant was aspirated and discarded, then washed with 70% ethanol by centrifugation for 15’ at 15 kRCF at room temperature. The supernatant was again aspirated and discarded, then the pellets were allowed to air dry for 30’ and finally resuspended in milliQ water. Resuspended samples were shaken for at least 1 hr at 37 °C for full resuspension.

### Characterizing cssDNA products from bacterial cssDNA preparations

The concentration of DNA produced by eScaf was quantified by UV-vis spectroscopy (NanoVue Plus). The purity of DNA products was visualized by agarose gel electrophoresis. All eScaf-produced DNA were run on 2% agarose in 0.5x TBE with 10 mM MgCl_2_ and prestained with SYBR-Safe. Gels were run for 2 hours at 100 V on ice using 0.5x TBE with 10 mM MgCl_2_ as running buffer. Gels were scanned using a Typhoon biomolecular imager (Cytiva). DNA ladders were the 1 kb DNA Ladder from New England Biolabs (NEB). Reference lanes contained cssDNA of the indicated length.

On-target purity was estimated by gel densitometry. DNA products from each preparation were run on separate lanes. To calculate each lane estimate, a rectangular region of the lane was selected from the gel image. Intensity of pixels within the region was background-subtracted, then averaged across the length of the lane to calculate a 1D line intensity average. A region corresponding to the band was selected and the area under the curve for that region was calculated. The percent purity was calculated by taking the area under the curve divided by the total integral of the 1D intensity average.

### Measuring eScaf growth in stirred-tank bioreactors

The bioreactor experiments used the original eScaf strain: *E. coli* TG1 with a single, untagged M13mp18 genome integrated at ulaD. Cultures were grown in 250 mL stirred-tank bioreactors (Culture Biosciences), with pH controlled at 7 and dissolved oxygen (DO) at 30%. Briefly, 50 mL eScaf seed cultures were grown overnight using LB with antibiotics in 250 mL shake flasks. After 24 hours, 3∼5 mL of seed culture was used to inoculate 150 mL eScaf media in 250 mL stirred-tank bioreactors (target OD after inoculation: 0.1) for the batch phase. Once the DO level spiked, indicating that the carbon source had been exhausted, feeds were turned on. Two different feed media were used, one containing glycerol (600 g/L glycerol,16 g/L tryptone, 10 g/L yeast extract, 50 µg/mL spectinomycin) and one containing glucose (750 g/L glucose, 20 g/L MgSO_4_, 18.5 g/L (NH_4_)_2_SO_4_, 50 µg/mL Spectinomycin). The cultures were either fed using a fixed glycerol feeding strategy or a dynamic glycerol or glucose feeding strategy. For cultures with a fixed glycerol feed, the first 7 hours of growth used an exponential feed, followed by a constant feed for the rest of the growth period. For cultures with a dynamic feed, the feeds were delivered as a variable-size bolus roughly every 30 minutes. During growth, 2 mL aliquots were taken at 12 different time points for CEDEX measurements and cssDNA purification. The pH of the cultures was controlled by addition of 14% ammonium hydroxide, and all cultures contained 10% polypropylene glycol 2000 as an antifoaming agent.

At the end of the growth period, the cultures were cooled to 10 °C, then pelleted at 3428 RCF, 4 °C for 30 minutes. The supernatant containing the phage particles was separated from the pellet and cssDNA was extracted from the phage and characterized as described above.

### Constructing His-tagged M13 donors and generating eScaf variants

eScaf variants were constructed via the INTEGRATE method (13). Plasmids for the INTEGRATE system were obtained from Addgene (pSL1119, pSL1777, Sternberg Lab). Target guide RNA sequences were cloned into “pEffector” (pSL1777) via site directed mutagenesis. The M13mp18 donor was cloned into “pDonor” (pSL1119) using golden gate cloning. His-tagged versions of the M13mp18 donor were generated by first adding an N-terminus 6xHis tag to the gene of interest via site-directed mutagenesis, then assembling the mutated M13mp18 cassette into the pSL1119 backbone by golden gate cloning.

Integration was performed by co-transforming the effector plasmid and donor plasmids into the host strain and incubating at 30 °C for 24∼48 hours. Transformants were screened for proper integration using cPCR. Positive transformants were grown overnight at 30 °C in LB with no antibiotics to cure the effector and donor plasmids. The overnight culture was plated on LB agar and grown overnight at 30 °C. The next day, colonies were patch plated onto an LB agar plate, then replica plated onto selection plates to ensure the plasmids had been fully cured. Fully cured colonies (no growth on either selection plate) were then grown in LB again for banking and to create competent cells.

Chemically competent eScaf cells were made by growing eScaf cultures in LB at 30 °C with shaking until the OD600 reached 0.5∼0.6. The cultures were put on ice for 10’, then pelleted at 3000 RCF, 4 °C for 15’. The supernatant was removed and the cells were resuspended in transformation storage solution (LB with 10% w/v PEG-3350, 5% v/v DMSO, 20 mM MgCl_2_), aliquoted, and either immediately used for transformation or stored at -80 °C.

### Replication and analysis

Replicates were independent biological cultures. Bars show means and error bars show one standard deviation where replicate measurements were available. Sample sizes are given in the figure legends.

## Results

### Culture medium affects eScaf growth, yield, and purity

As a first step to increase cssDNA yield, we grew eScaf in media that were richer in nutrients than lysogeny broth (LB). We transformed a phagemid encoding a 3031-nt cssDNA sequence into eScaf and grew the transformants in LB, 2x Yeast Extract Tryptone media (2xYT), or Terrific Broth (TB) and harvested the cssDNA (Fig 1a). We found the resulting DNA yield from cultures grown in TB, the most nutrient dense of the three media, to be highest at 0.525 mg per liter of culture, followed by yield from 2xYT-grown cultures at 0.29 mg/L, and then from LB-grown cultures at 0.122 mg/L, which is consistent with our previous work. However, we also found that DNA from TB cultures contained more off-target material than DNA harvested from 2xYT or LB cultures, which had little-to-no off-target production. On average, DNA from TB cultures were about 87.4% pure, whereas DNA from the LB and 2xYT cultures were about 93.5% and 95% pure, respectively. From these data, we estimate that on-target cssDNA yield of cultures grown in TB was still roughly 0.46 mg/L, substantially higher than full DNA yields from either LB or 2xYT-grown cultures.

**Figure 1.**
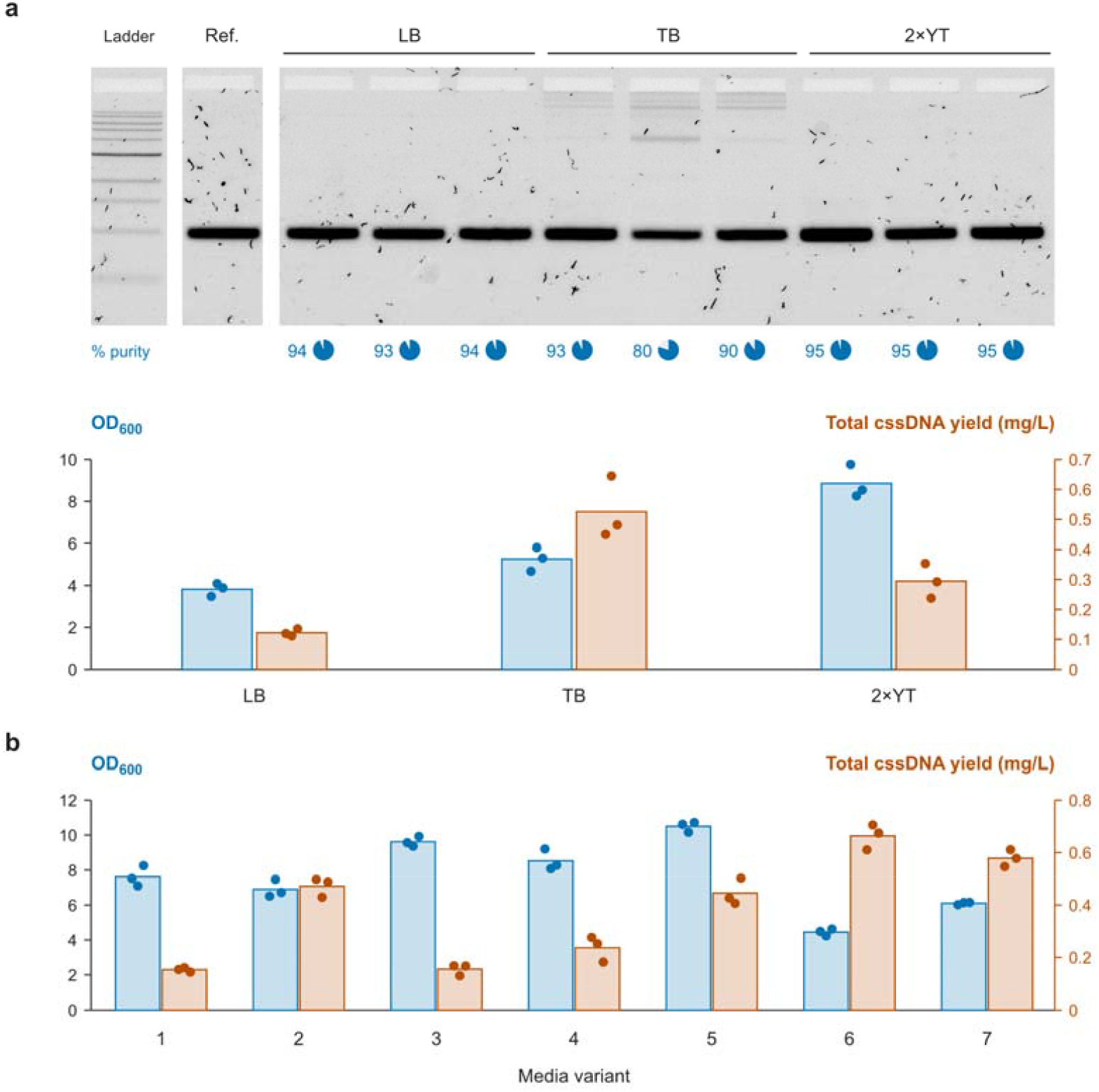
Media optimization of cssDNA production by eScaf. (a) Comparison of total cssDNA yield, on-target purity, and endpoint cell density for eScaf grown in LB, TB, or 2×YT (n = 3 biological replicates). TB cultures produced the highest total cssDNA yield (0.53 mg/L) but had the lowest mean on-target purity (87%). Cultures grown in 2×YT yielded 0.29 mg/L with 95% mean on-target purity. Blue and orange bars show endpoint cell density and total cssDNA yield, respectively; purity values beneath the gels refer to individual lanes. (b) Total cssDNA yield and endpoint cell density across seven custom complex media variants (Table 1; n = 3 biological replicates per condition). Medium #6 (eScaf medium, EM) produced the highest total cssDNA yield (0.66 mg/L), with 95% mean on-target purity, and the lowest endpoint cell density among these variants. Bars show means; points show individual measurements. Ref., cssDNA reference of the indicated length. Ladders, NEB 1 kb DNA Ladder.

We found that using 2xYT resulted in the most eScaf growth compared to using LB or TB. Specifically, we observed that the growth of the 2xYT cultures, as measured by OD600, was more than 2x the growth of the LB cultures, commensurate with the increase in cssDNA yield. However, the cells grown in TB only reached a final OD600 about 1.35x higher than that of the LB culture, but the on-target cssDNA yield was more than 3x as high. Thus, the low growth relative to the on-target cssDNA yield suggests that TB enhances average production of M13 particles per individual cells relative to LB. 2xYT does not appear to enhance eScaf productivity, but does appear to promote growth compared to LB, leading to a higher cssDNA yield per unit volume of culture.

**Table 1.** Complex media recipes.

| Medium | Tryptone<br>(g/L) | Yeast extract<br>(g/L) | NaCl<br>(g/L) | KH<sub>2</sub>PO<sub>4</sub><br>(g/L) | K<sub>2</sub>HPO<sub>4</sub><br>(g/L) | Glycerol<br>(mL/L) |
| --- | --- | --- | --- | --- | --- | --- |
| 2xYT | 16 | 10 | 5 | 0 | 0 | 0 |
| TB | 12 | 24 | 0 | 2 | 13 | 4 |
| 1 | 16 | 10 | 5 | 2 | 13 | 0 |
| 2 | 16 | 10 | 5 | 0 | 0 | 4 |
| 3 | 16 | 10 | 5 | 2 | 13 | 4 |
| 4 | 16 | 24 | 5 | 0 | 0 | 4 |
| 5 | 20 | 10 | 5 | 0 | 0 | 4 |
| 6 | 16 | 10 | 0 | 2 | 13 | 4 |
| 7 | 16 | 24 | 0 | 2 | 13 | 4 |

We then aimed to develop a media formulation that could achieve the same or greater productivity as TB while maintaining the purity achieved in 2xYT. We created seven different media variants based on 2xYT and TB. We screened the cssDNA produced by eScaf grown in these media variants (Table 1, Fig 1b).

We obtained the highest yield and best purity from media #6, which we call “eScaf media” (EM). EM results in the lowest eScaf endpoint cell density of all media we tested other than LB, suggesting that EM substantially favors productivity over growth. EM has the same concentration of tryptone and yeast extract as 2xYT but does not include NaCl and includes glycerol and phosphate buffer. If NaCl is included in the mixture (Media #3) we find that growth more than doubles, but yield decreases by over a factor of 4, resulting in the lowest productivity culture. We found that media containing both NaCl and phosphate (media #1, media #3) were the least productive.

Media #7 was the second most productive. Productivity was similar to TB but also contained off-target species. Media #7 had an average purity of 83%, compared to the 95% purity we achieve with EM. Media #7 has the same concentration of tryptone as 2xYT (16 g/L) but the same concentration of yeast extract as TB (24 g/L) and shares a higher percentage of off-target DNA production similar to TB. eScaf grew faster in media #7 than in TB, suggesting that the increased tryptone facilitates growth.

Medium #5, which contains additional tryptone, supported the highest endpoint cell density but produced less cssDNA than eScaf medium. Medium #4, which contains additional yeast extract, supported lower cell density and yield than medium #5.

We note that in terms of improving overall yield, adding an extra carbon source in the form of glycerol (Media #2) dramatically improved cssDNA yield compared to 2xYT. Simply adding glycerol to 2xYT increased cssDNA yield to 0.472 mg/L. Taken together, we optimized a complex media for eScaf that enables us to harvest, on average, 0.663 mg of cssDNA per liter of culture with high purity. Relative to cssDNA yields grown by eScaf in LB, this represents an increase of nearly 5.4x. At this yield, a single 2 pmol dose of a 3000-nt repair template for therapeutic CRISPR knock-in would be achievable from a 3 mL culture, enabling high-throughput screening assays in 24-well plates.

### eScaf grows poorly in M9 and Riesenberg media

To minimize prep-to-prep cssDNA variability and gain finer control over media composition, we experimented with growing eScaf in chemically defined media by replacing variable components such as tryptone and yeast extract with a specific chemical cocktail. We grew eScaf in shake flasks at 30 °C with M9 minimal media supplemented with 2% glucose, 0.1 mM CaCl2, 1 mM MgSO4, and 0.5 µg/mL thiamine and observed very slow growth. We allowed cultures to continue growing beyond the typical 16-hour endpoint and harvested after 40 hours of growth. The mean OD600 of harvested cultures was only 0.88±0.13, with mean yield of 3.98±0.95 µg of DNA per liter of culture (SFig. 1a). To attempt to produce eScaf at a higher rate, we grew eScaf in shake flasks at 30 °C in Riesenberg media (14, 15) as well as in 13 other Riesenberg media variants. As before, we extended growth time beyond 16 hours and harvested after 24 hours of growth. We found that eScaf grown in some Riesenberg media variants could reach a final concentration similar to that of a culture grown in complex media, but we could not get sufficient cssDNA yield or quality in Riesenberg media or any variant we tested so we did not pursue further optimization of defined media (SFig 1b).

### Closed-loop production of cssDNA in bioreactors reaches mg/L scale

Closed-loop control over microbial growth in bioreactors enables far higher cell densities and is key to increasing scalability of bioproduction. We grew eScaf cultures in 250 mL bioreactors over 67 hours using three different feed strategies. In the first strategy, we fed in a glycerol-based media continuously, starting with an exponentially increasing amount and then leveling off to a constant amount (“exp-to-constant feed”). In the second strategy, we fed in the same glycerol-based media but introduced a bolus at a specific time (“spike feed”). In the third strategy, we used a glucose-based media and spike-fed it to cells in the same manner as the second feed strategy. All cultures were grown in eScaf media. Culture aliquots were collected at various points during the 67-hour growth period and processed to measure metrics such as OD583, cssDNA yield, and cssDNA purity over time. These data are plotted in Fig. 2a–c.

**Figure 2.**
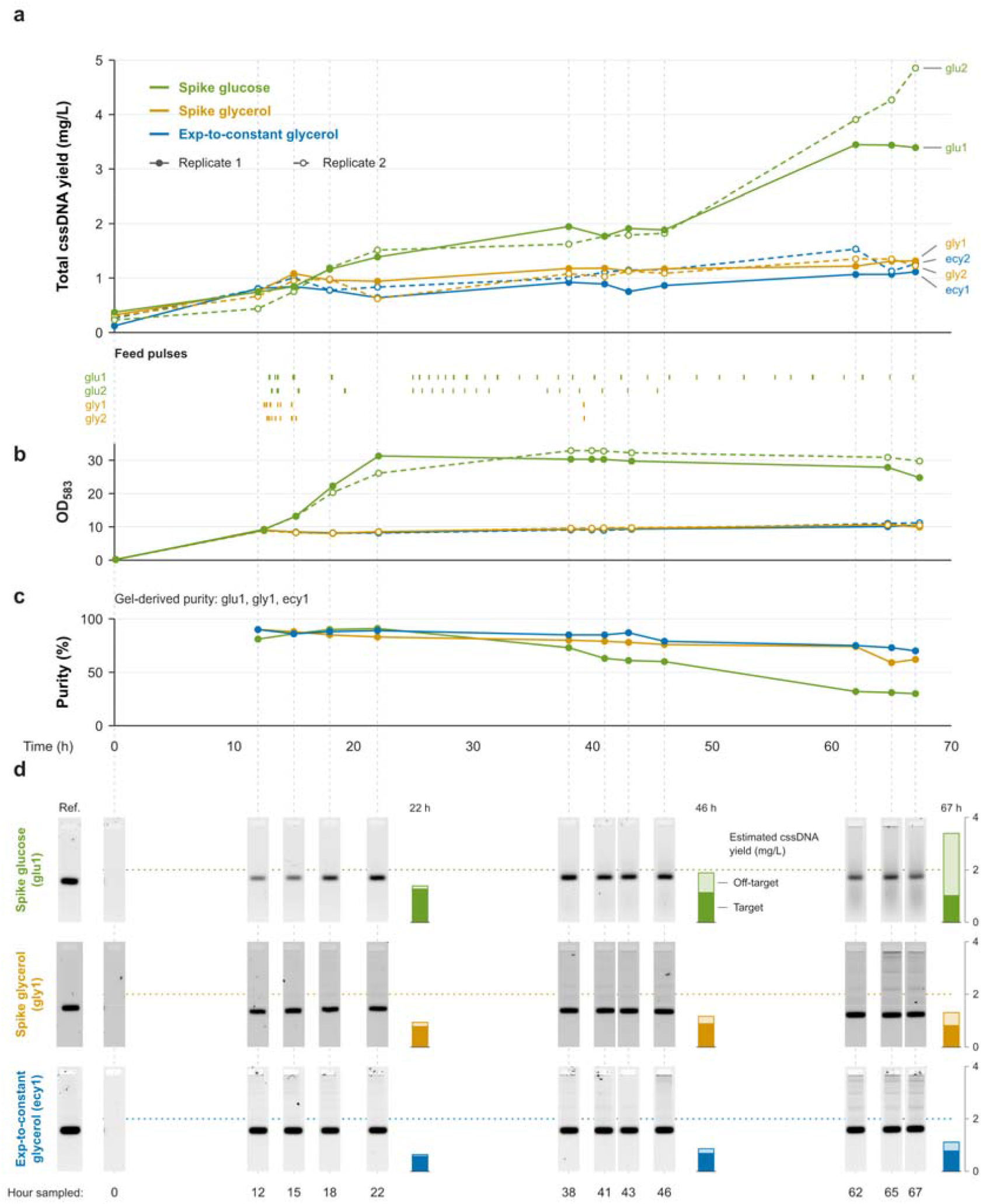
eScaf growth and DNA production in stirred-tank bioreactors. (a) Total cssDNA yield and (b) cell density over 67 hours of bioreactor growth under three feeding strategies, with two biological replicates per condition. Glucose spike-fed cultures reached the highest endpoint yield (4.12 mg/L) and cell density (OD583 ∼25–30). Spike and exp-to-constant glycerol-fed cultures yielded 1.27 and 1.19 mg/L, respectively, and reached OD583 ∼10. Feed annotations show recorded pump activity. (c) On-target cssDNA purity over time, estimated by gel densitometry. Glucose spike-fed cultures reached their highest purity at 22 hours (91%), declining to 30% by 67 hours. Endpoint purities for spike and exp-to-constant glycerol feeding were 62% and 70%, respectively. (d) Gel analysis of samples from one biological replicate per feeding condition (glu1, gly1, and ecy1). Stacked bars at 22, 46, and 67 hours show estimated on-target and off-target cssDNA yields, calculated from the corresponding total cssDNA yield and gel-derived purity. Estimated target cssDNA yield = total cssDNA yield × gel-derived target fraction; the remaining bar segment is the difference. Lanes within each feeding condition were cropped from the same gel; different feeding conditions were run on separate gels. Ref., cssDNA reference of the indicated length.

At 67 hours, the time course samples from glucose spike-fed cultures had the highest mean total cssDNA yield, 4.12 mg/L. Spike and exp-to-constant glycerol-fed cultures yielded 1.27 and 1.19 mg/L, respectively (Fig. 2a). Gel analysis showed more background smearing in the glucose-fed culture and a stronger ladder pattern in the glycerol-fed cultures. Endpoint on-target purity was 30% for glucose spike feed, 62% for glycerol spike feed, and 70% for exp-to-constant glycerol feed (Fig. 2c,d).

Examining the cssDNA quality over time, we observed that the target is the dominant product in the glycerol cultures. For cultures given exp-to-constant glycerol feeds, production of the target band stabilizes by 12 hours, with the highest purity at 90%. Subsequently, off-target DNA production increases. Cultures with spike glycerol feeds showed maximum cssDNA production at the 15-hour time point, with increasing off-target DNA production after that time point. Cultures with spike glucose feeds, in contrast, produced off-target DNA, with only 60% on-target products by 46 hours of growth and 30% on-target by the end of the 67-hour growth period. Glucose spike-fed cultures reached their highest on-target purity at 22 hours (91%), after which off-target products increasingly accumulated.

Comparing cssDNA yield and growth over time among media conditions, we observed that glucose-fed eScaf cultures saturate at OD583 between 25 and 30 compared to glycerol-fed eScaf cultures, which saturate at closer to an OD583 of 10 for both feed strategies (Fig. 2b). Although both glycerol-fed cultures and glucose fed cultures grew at similar rates for the first 12 hours, afterwards, the glycerol-fed culture growth hit its maximum and began leveling off. However, the glucose-fed cultures continued to grow, reaching stationary phase roughly 22 hours in. The time series data show that eScaf begins off-target species production roughly around the same time as reaching stationary phase. At 12 hours, mean total cssDNA yields were 0.8 mg/L for exp-to-constant glycerol feeding and 0.7 mg/L for spike glycerol feeding. Glucose spike-fed cultures reached a mean total cssDNA yield of 1.5 mg/L at 22 hours.

### Host strain genotype affects eScaf cssDNA quality

Previously, we used TG1 as the original eScaf host strain due to its historical usage as a phage-display strain. However, since eScaf does not rely on M13 infection, we do not require our host strain to have an F’ episome. Therefore, we are not restricted in choice of potential host strains into which M13 can be integrated. Given that some host strains have been engineered for high yield plasmid production or for cloning difficult sequences, we reasoned that integrating the M13 genome into alternate host strains could be a useful way to improve cssDNA quality. We screened two commercially available *E. coli* strains, 5α and 10β, both from New England Biolabs, to assess whether host genotype affects cssDNA yield.

We found substantial differences in yield among the three different eScaf strains. The eScaf strain derived from 5α (eScaf-5α) had the highest yield, more than 2x that of our original eScaf strain (Fig. 3a). The eScaf strain derived from 10β (eScaf-10β) had the lowest yield, below 0.5x that of our original eScaf strain. The cell densities at saturation between eScaf-5α and the original eScaf cultures were comparable (Fig. 3a), revealing that the eScaf-5α strain is more productive than the original eScaf. In contrast, the final cell density of eScaf-10β was substantially lower, showing eScaf-10β has the lowest productivity of the three. Additionally, we noted that the DNA from eScaf-10β contained more off-target products than the original eScaf or eScaf-5α (Fig. 3b). Thus, we find that the host genotype affects eScaf cssDNA quality, with eScaf-5α having the highest yield.

**Figure 3.**
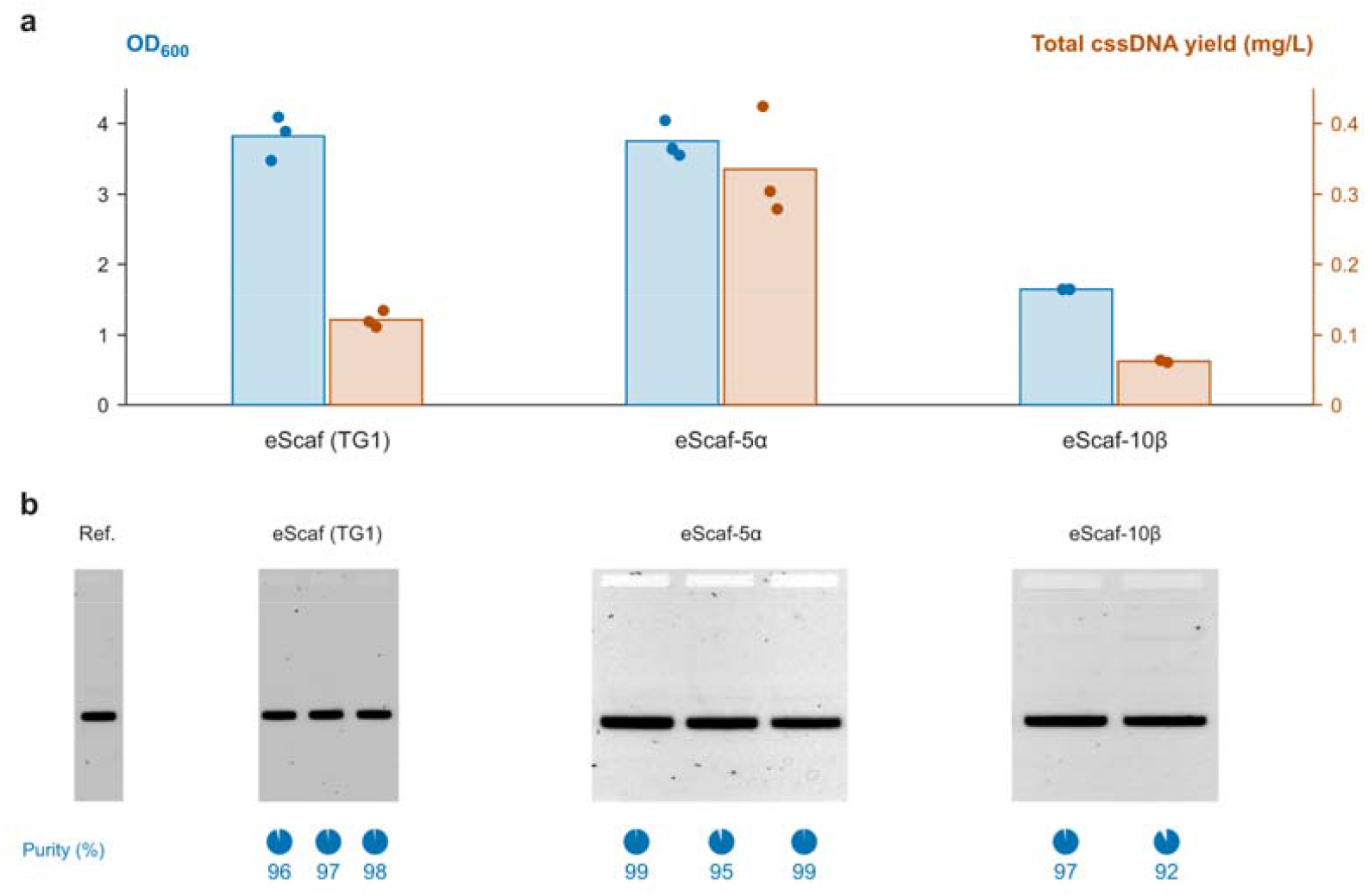
Effect of host strain on cssDNA production. (a) Total cssDNA yield and endpoint cell density of eScaf strains constructed by integrating the M13 genome into TG1, NEB 5α, or NEB 10β (n = 3 biological replicates for TG1 and 5α; n = 2 for 10β). eScaf-5α produced more than twice the yield of TG1-based eScaf at comparable cell density. eScaf-10β had the lowest yield and cell density. (b) Gel analysis of cssDNA products from each host strain, with on-target purity estimated by densitometry. A uniform gamma adjustment was applied to the eScaf-5α gel for display; purity was quantified from the unadjusted source data. Ref., cssDNA reference of length 3031 nt.

### His-tagging g3 increases eScaf productivity

To facilitate processing, we added His-tags to the phage coat (16, 17). We cloned three different transposon donors carrying the M13mp18 genome with a 6xHis tag on the N-terminus of either pIII, pVII, or pVIII. To confirm the edited M13mp18 genome was working properly, we tested cssDNA production using the His-tagged M13 expressed from the donor as a helper plasmid in TG1.

We found that, while our His-tagged pIII and pVII donor plasmids produced the target cssDNA, the His-tagged pVIII donor plasmid was unable to make the target product (Fig. 4a). The cssDNA from the His-tagged pIII donor was clean, but the cssDNA from the His-tagged pVII donor had a clear off-target band. We found that the His-tagged pIII donor produced nearly 6x the yield of the His-tagged pVII donor and 10x the yield of original eScaf strain, with a small shake flask culture reaching more than 1.5 mg of cssDNA per liter of bacterial culture.

**Figure 4.**
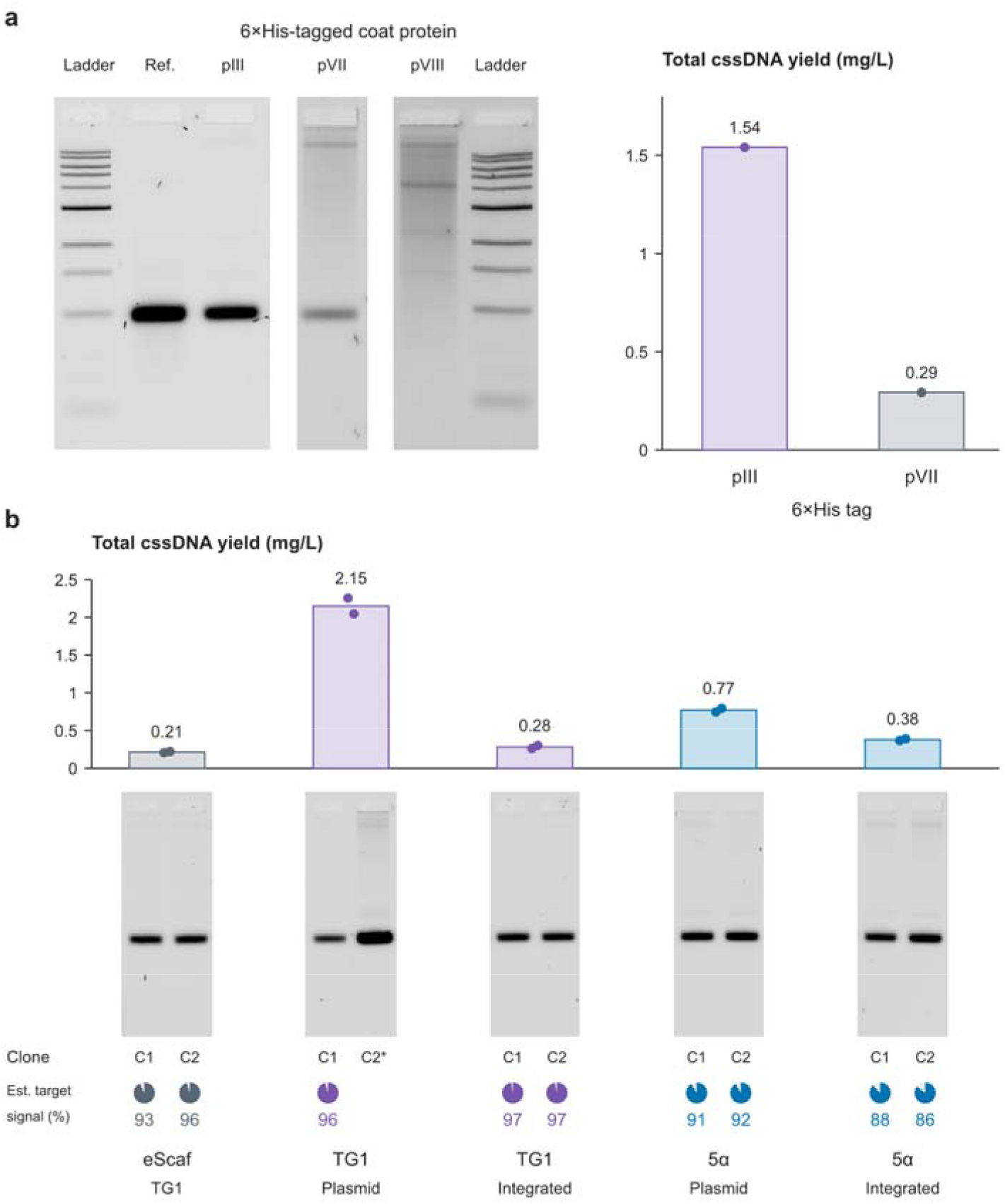
Effect of His-tagging M13 coat proteins on cssDNA production. (a) Gel analysis and total cssDNA yield from TG1 carrying M13 helper plasmids encoding His-tagged pIII, pVII, or pVIII. His-tagged pIII yielded 1.5 mg/L, whereas His-tagged pVII yielded 0.29 mg/L and produced off-target species (n = 1 per quantified condition). No target cssDNA band was detected with His-tagged pVIII. (b) Total cssDNA yield from the original eScaf strain and from TG1 or NEB 5α carrying the His-tagged pIII M13 helper on a plasmid or integrated into the chromosome (n = 2 biological replicates per condition). Plasmid-based helpers produced higher yields than integrated helpers in both hosts. With plasmid-based helpers, TG1 yielded approximately 2.8-fold more cssDNA than 5α. Gels show clones C1 and C2 for each condition. Lanes were loaded with 100 ng DNA, except TG1 plasmid C2, which contained approximately 350 ng. On-target purity was estimated by densitometry; the overloaded C2 lane was excluded. Ref., cssDNA reference of length 3031-nt. Ladders, NEB 1 kb DNA Ladder.

We integrated the His-tagged pIII M13 into both TG1 and NEB 5α and measured total cssDNA yield (Fig. 4b). Again, we saw high yields when producing cssDNA with the His-tagged pIII donor as a helper plasmid. Producing cssDNA with NEB 5α expressing the His-tagged pIII donor gave high yields, but not nearly as high as the TG1-based system. We found that integrating the His-tagged donor resulted in lower total cssDNA yield than when producing cssDNA on plasmid. We found that the integrated version of 5α again produced higher yields than the integrated version of TG1. Circular ssDNA production using the His-tagged pIII helper plasmid in TG1 had substantially higher yield than in 5α, the only condition in which the TG1 host strain outperformed the 5α strain. The mechanism for this difference is still unclear.

### Increasing copy number increases eScaf yield

Based on the His-tagging results, we investigated the effect of M13 copy number on cssDNA production. Previously, we proposed that by reducing the M13 genome copy number in *E. coli* to a single copy, we could minimize M13 toxicity which would, in turn, result in high-quality cssDNA production. This resulted in the high purity eScaf strain, albeit with low yield. Our yield differences between the episomally expressed His-tagged helpers and the chromosomally integrated His-tagged helpers suggest that higher copy number increases DNA yield and given that the cssDNA from the His-tagged g3 helper plasmid was relatively pure, high copy number M13 expression could be achieved without severe *E. coli* toxicity. We reasoned that there could be an optimal copy number for M13 that balances toxicity and productivity.

To test the effect of M13 genome copy number on cssDNA yield, we added a second copy of the M13mp18 genome to eScaf. To determine where to integrate the second M13mp18 genome, we screened integration sites to see how they affected cssDNA yield (Fig. 5a, Table 2). We fixed one integration site to the highest performing loci (Site #6). The second locus was either at a high-yield site (Site #4) or a low-yield site (Site #2). We found that the double-integrated strains had triple the yield of the original strain, even for the variant with a low-yield second integration site. In line with our single-insertion site screen, the double integrated variant in which we integrated the second highest yield site had higher overall yield than the double integrated variant with the second lowest yield site. We tested double integration using the His-tagged M13mp18 donor in high yield sites 4 and 6. In agreement with our previous results, we found that the His-tagged variant of the double integrated strains had the highest yield (Fig. 5a).

**Table 2.**
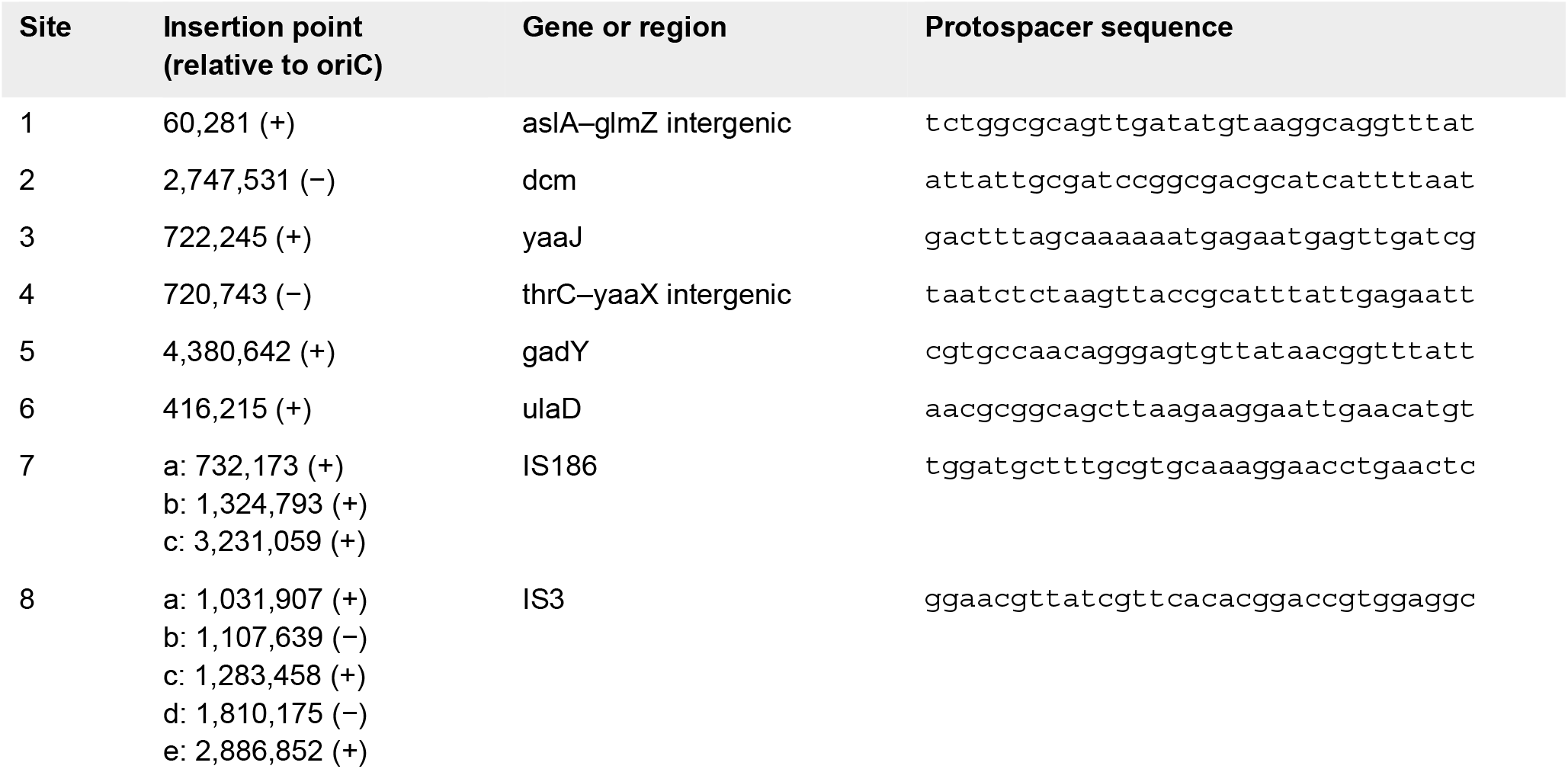
Integration sites and protospacer sequences.

**Figure 5.**
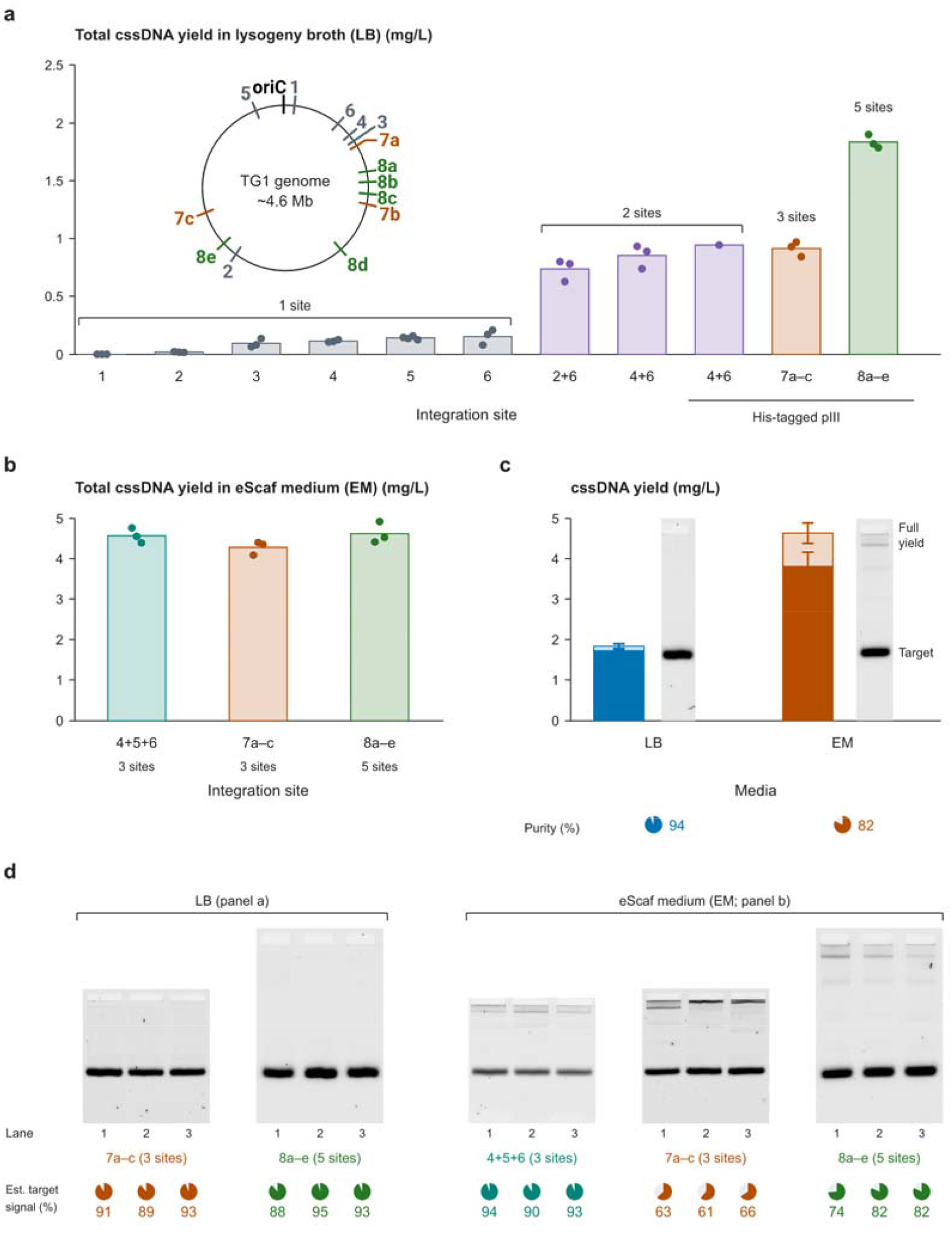
M13 copy number and integration location affect cssDNA yield. (a) Total cssDNA yield in LB for strains with the indicated M13 integration sites and copy numbers. eScaf-IS3, with five integrated copies, produced the highest yield. Biological replicates: n = 3 per condition, except site 5 (n = 4) and His-tagged 4+6 (n = 1). (b) Total cssDNA yield in eScaf medium (EM) for strains with three or five integrated M13 copies (n = 3 biological replicates per condition). The strain with integrations at sites 4, 5, and 6 and eScaf-IS186 produced similar yields to the five-copy eScaf-IS3 strain. (c) Total and on-target cssDNA yields from eScaf-IS3 grown in LB or EM (n = 3 biological replicates per condition). EM supported higher total cssDNA yield but lower on-target purity than LB. On-target yields were calculated for each replicate using its gel-derived purity and then averaged. Error bars show one standard deviation. (d) Gel analysis of eScaf-IS186 and eScaf-IS3 grown in LB or EM, and the strain with integrations at sites 4, 5, and 6 grown in EM. On-target purity was estimated by densitometry. A uniform gamma adjustment was applied to the sites 4, 5, and 6 gel for display; purity was quantified from the unadjusted source image. Ref., cssDNA reference of the length 3031 nt.

### DNA yield depends on both copy number and integration site

We wondered if cssDNA yield would continue to increase with copy number or if there is an inflection point after which increased M13 expression causes cssDNA quality to degrade. To test this idea, we designed three different gRNA sequences targeting different insertion sequences (IS). We used multi-copy sequences in the *E. coli* genome (18) that have previously been shown to accommodate multiple parallel insertions. We targeted IS186, IS3, and IS1, which have 3, 5, and 7 copies in TG1, respectively. We found that we were unable to integrate the M13 donor into IS1, but we were able to integrate into IS3 (eScaf-IS3) and IS186 (eScaf-IS186). cPCR results confirmed that all IS3 and IS186 loci were properly integrated. To test if 6 copies of the M13 genome could be integrated into TG1, we made eScaf-IS3 competent and attempted to integrate one more copy into eScaf-IS3 at insertion site #6, but we observed no colonies after integration.

In LB, eScaf-IS3, which contains five integrated M13 copies, produced more total cssDNA than eScaf-IS186, which contains three copies (Fig. 5a). However, the eScaf-IS186 variant had very similar yield to the double-integrated version despite having one more copy of the His-tagged M13mp18 genome integrated into its genome (Fig. 5a). We reasoned that the similar yields between eScaf-IS186 and the double-integrated eScaf variant could be due to integration location. To test this, we made another variant of eScaf with three copies of the His-tagged M13 helper at the three strongest single integration sites (integration sites #6, #5, #4) and compared yields in eScaf medium (EM). In EM, both three-site variants produced similar total cssDNA yields to the five-site eScaf-IS3 strain (Fig. 5b), unlike the higher yield of eScaf-IS3 observed in LB. Thus, our results suggest that both copy number and integration location play a role in overall cssDNA yield.

Currently, eScaf-IS3 represents the highest yield eScaf strain we have engineered so far. In LB media we produced close to 2 mg/L of cssDNA. In eScaf medium, we produced an average total cssDNA yield of 4.6 mg/L, with an average on-target purity of 82% (Fig. 5c). The estimated on-target cssDNA yield, calculated for each replicate and then averaged, was 3.8 mg/L.

## Discussion

Our improvements in yields were substantial but could be improved still further with additional bioreactor optimizations. The optimizations described above enhanced cssDNA yields in shake flasks relative to our original eScaf preparations by nearly 40x: from ∼100 µg/L to ∼4 mg/L. Our current yield is comparable to typical helper plasmid-phagemid systems grown in bioreactors or flasks as reported in the literature (5). We tested bioreactor feeding strategies using a single-copy eScaf strain. Applying these strategies to a multicopy variant such as eScaf-IS3, together with further optimization of fermentation conditions, could increase cssDNA yield. Careful screening of bioreactor conditions for cssDNA production using wild-type phage, helper phage, or “artificial” phage in bioreactors has resulted in cssDNA yields in the 100s of mg/L (8, 9, 19), indicating substantial room for optimizing growth in bioreactors.

The yields our work enable can open doors to entirely new applications of cssDNA. The yield improvements we described here may enable applications that were previously gated by the difficulty of producing cssDNA. In cellular engineering, for example, cssDNA has been shown to be a potent donor for homology-driven repair, with cssDNA constructs used to successfully knock-in payloads > 5 kb with up to 40% efficiency and minimal toxicity (20). The scale of cssDNA produced by eScaf-IS3 now allows for producing and screening cssDNA donors at high-throughput, either by testing many constructs in parallel or conducting pooled screens across many conditions. Our custom cssDNA production system feeds naturally into structural DNA nanotechnology applications, and the scale and length of our cssDNA open the door for much larger DNA origami monomers with programmable scaffold sequences for designing even larger multi-origami constructs. Finally, several methods exist to excise ssDNA fragments from the larger cssDNA product (21, 19). Using these methods, eScaf could become an excellent method to amplify ssDNA fragments by cloning those sequences into a phagemid, producing cssDNA, and then releasing the desired oligos from the cssDNA and gel purifying them. This technique would offer many benefits ranging from producing aptamers to DNA displacement circuits.

### Our results suggest that we could further manipulate the host genome with positive results

Examining differences between the NEB 5α and TG1 genomes could reveal which genes promote cssDNA production. For example, the increased yield in eScaf-5α could be related to 5α having disabling mutations of endonuclease I and recA, whereas TG1 has functional copies of both proteins. Since the cssDNA produced by eScaf is templated off a phagemid, endonuclease I activity resulting in degradation or recA activity resulting in recombination of the phagemid could result in lower cssDNA yield. Alternatively, because TG1 is an F’ strain, maintenance of the F episome could divert energy from producing phage particles; given that the eScaf-TG1 and eScaf-5α ODs were similar at saturation, it is reasonable to assume that the metabolic burden from F episome replication could result in lower cssDNA yield. Future work should use transcriptomic and metabolomic analyses to inform genome edits that could significantly boost yield.

We introduced the His-tag on the M13 pIII coat protein to provide handles for column purification or attachment of fluorescent tags for visualization. We found that His-tagging pIII also increased cssDNA yield relative to wildtype pIII, an effect that merits further investigation. In the natural M13 lifecycle, pIII complexes with the *E. coli* pilus to initiate infection. In eScaf, we do not need infection to grow phage, and we hypothesized that the presence of the His tag could be interfering with the pIII-pilus complex which would ordinarily cause M13 particles to adsorb onto *E. coli* pili. This logic implies that His-tagged helpers yield more recoverable phage because particles that would otherwise adsorb to pili and co-pellet with bacteria are instead released into the supernatant. However, we saw a positive effect of His-tagged pIII even in the pilus-free strain NEB 5α, suggesting that our hypothesized “adsorption” mechanism was not correct. Considering these data, we propose an alternative hypothesis that the His-tag on the N terminus somehow facilitates export of the fully formed M13 particle, since removal of g3p does abolish M13 secretion. The C-terminus of pIII is the necessary domain for secretion, but it is not impossible that the N-terminus His modification would influence M13 export in some way. (22).

The higher yield from plasmid-based helpers compared with integrated helpers could reflect differences in M13 copy number. Among integrated-helper strains, 5α outperformed TG1, consistent with our earlier host-strain comparison. With plasmid-based helpers, TG1 produced approximately 2.8-fold more DNA than 5α, the only condition in which TG1 outperformed 5α. It seems likely that the mechanism which enhances His-tagged pIII M13 export could still be related to the TG1 pilus and when M13 is expressed episomally with high copy number, the effect of secretion enhancement is much greater on TG1 than the F-5α strain. Further study of how N-terminal pIII modifications affect M13 export could inform improvements in cssDNA production and phage display, as well as our understanding of M13 biology.

The cssDNA yields we report here are for a single sequence and different sequences may return different results. One limitation of this work is that our metrics are based on a single sequence, and we have observed that changing sequence can cause significant variation in yield. Fully determining how sequence variation affects yield was beyond the scope of our investigations. In this work, we used a 3031-base sequence derived from the coding section of human dynein (23). We compared this sequence to three other 3000-base sequences and found that its yield was closest to the average of yield of the four sequences. Thus, here, we present what we consider to be an average yield. The exact yield of a specific, user-defined, sequence will need to be determined empirically. The optimizations described here should still hold regardless of the sequence identity.

Future work. Medium composition affected both cssDNA yield and product purity. TB and medium #7, both rich in yeast extract, produced more off-target material than eScaf medium. Further medium optimization could help improve yield while limiting off-target production. For example, Kick *et al*. observed that (NH_4_)_2_HPO_4_ produces fewer background products than K_2_HPO_4_ (8). Iterative and high-throughput screening of integration locations in eScaf could determine the optimal copy number and location of integrated M13 genomes (24). Other high-throughput genomic tools, such as CRISPRi (25) and CRISPRa (26) could help identify and modulate genes that improve phage production. Additionally, further modifying the M13 genome or the pScaf phagemid backbone could increase cssDNA yield and quality (27, 28).

## Conclusion

By optimizing media composition, host strain, and M13 genome copy number and integration location, we increased eScaf cssDNA yield in shake flasks by nearly 40-fold, from ∼100 µg/L to ∼4 mg/L. Bioreactor experiments with a single-copy eScaf strain identified feeding strategies that could support further improvements. The resulting eScaf-IS3 strain now produces cssDNA at yields competitive with helper plasmid-phagemid systems, crossing practical thresholds for use in screening to find efficacious nucleic acid therapies, multi-operon-length gene donor production for engineered T cells, and large-scale DNA origami assembly for viral trapping or photonic crystals. The unexplained yield boost from His-tagged pIII suggests that there may be more ways to increase M13 yield based on the mechanics of the M13 lifecycle; yield variation due to host strain genotype points to additional engineering headroom. Finally, further bioreactor optimization suggests yields well above 4 mg/L are within reach. Our eScaf-IS3 system provides a robust, consistent foundation for both immediate cssDNA applications and continued scale-up toward commercial biomanufacturing.

## Acknowledgments

pSL1777 (pEffector) and pSL1119 (pDonor) were gifts from Samuel H. Sternberg (Addgene plasmid# 160731; http://n2t.net/addgene:160731; RRID:Addgene 160731, Addgene plasmid# 160732; http://n2t.net/addgene:160732; RRID:Addgene 160732). We thank Hannah Monroe, Kristina Tyler, Rebecca Du, and Cayley Hoyer of Culture Biosciences for discussion and guidance in media development and bioreactor processing. We thank Leslie Bienen of C3 Science Communication for editing and feedback on the manuscript.

During manuscript preparation, we used OpenAI Codex (GPT-6-Astra) to assist with language editing, figure formatting, and development of code for data processing and visualization. The authors reviewed and edited the resulting material and take responsibility for the final manuscript.

## Author contributions

KS and SMD designed experiments, analyzed data, and wrote the manuscript. KS performed all experiments.

## Funding

This work was supported by the National Institute of General Medical Sciences of the National Institutes of Health under Award Number R35GM125027. The content is solely the responsibility of the authors and does not necessarily represent the official views of the National Institutes of Health.

## Conflict of interest statement

None declared.

## Supplementary data

Supplementary information accompanies this preprint.

## Data availability

Data and code supporting this study are available at Zenodo (https://doi.org/10.5281/zenodo.23718450).

## Notes

https://zenodo.org/records/23718450

### Competing Interest Statement

The authors have declared no competing interest.

