## Supplementary Information for "Genomic and environmental determinants of circular single-stranded DNA production in engineered *E. coli*"

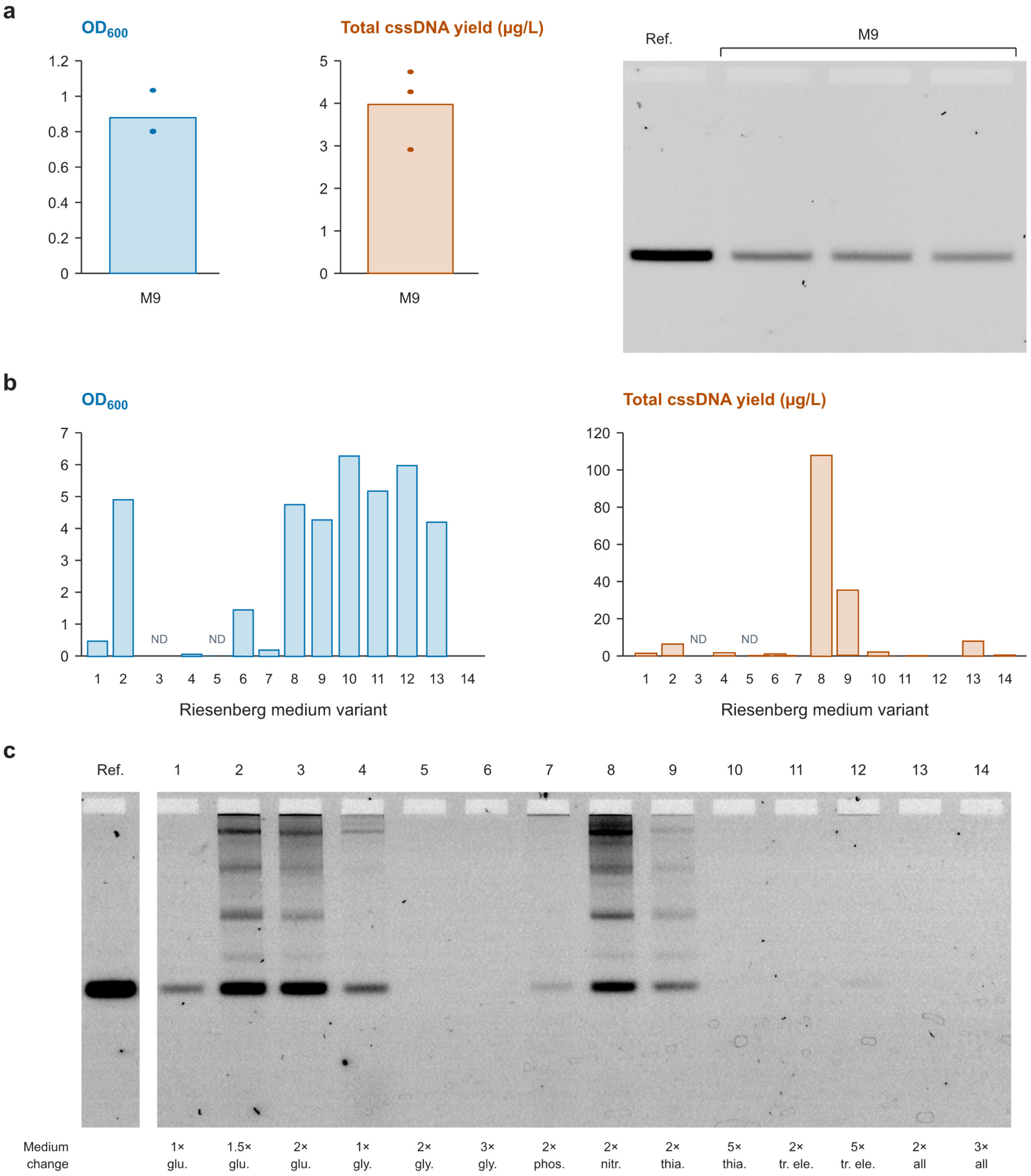


**Figure S1. eScaf growth and cssDNA production in M9 and Riesenberg media.** (a) Growth and total cssDNA yield of the original eScaf strain in M9 medium. Mean OD600 was 0.88 ± 0.13, and total cssDNA yield was 4.0 ± 1.0 µg/L (mean ± SD; n = 3 biological replicates). (b) Endpoint cell density and total cssDNA yield of the original eScaf strain grown in Riesenberg medium and its variants (n = 1 per condition). (c) Gel analysis of the resulting cssDNA products. The original Riesenberg formulation produced fewer off-target species than the tested variants, with a total cssDNA yield of approximately 1.4 µg/L. Ref.: 3031-nt cssDNA reference.

**Riesenberg media composition**

| Variant | $KH_2PO_4$ (g/L) | $(NH_4)_2PO_4$ (g/L) | Glucose (g/L) | Glycerol (mL/L) | EDTA (mg/L) | Thiamine (mg/L) | Citric acid (g/L) | $MgSO_4$ (g/L) | Trace elements |
|---|---|---|---|---|---|---|---|---|---|
| 1 | 13.3 | 4 | 20 | 0 | 8 | 4.5 | 1.7 | 1.2 | 1x |
| 2 | 13.3 | 4 | 30 | 0 | 8 | 4.5 | 1.7 | 1.2 | 1x |
| 3 | 13.3 | 4 | 40 | 0 | 8 | 4.5 | 1.7 | 1.2 | 1x |
| 4 | 13.3 | 4 | 0 | 4 | 8 | 4.5 | 1.7 | 1.2 | 1x |
| 5 | 13.3 | 4 | 0 | 8 | 8 | 4.5 | 1.7 | 1.2 | 1x |
| 6 | 13.3 | 4 | 0 | 12 | 8 | 4.5 | 1.7 | 1.2 | 1x |
| 7 | 26.6 | 4 | 20 | 0 | 8 | 4.5 | 1.7 | 1.2 | 1x |
| 8 | 13.3 | 8 | 20 | 0 | 8 | 4.5 | 1.7 | 1.2 | 1x |
| 9 | 13.3 | 4 | 20 | 0 | 8 | 9 | 1.7 | 1.2 | 1x |
| 10 | 13.3 | 4 | 0 | 12 | 8 | 22.5 | 1.7 | 1.2 | 1x |
| 11 | 13.3 | 4 | 0 | 12 | 8 | 4.5 | 1.7 | 1.2 | 2x |
| 12 | 13.3 | 4 | 0 | 12 | 8 | 4.5 | 1.7 | 1.2 | 5x |
| 13 | 26.6 | 8 | 40 | 0 | 16 | 9 | 3.4 | 2.4 | 2x |
| 14 | 39.9 | 12 | 60 | 0 | 24 | 13.5 | 5.1 | 3.6 | 3x |

**Trace element composition**

| Component | Concentration (mg/L) |
|---|---|
| CoCl2 · 6H2O | 4 |
| MnCl2 · 4H2O | 23.5 |
| CuCl2 · 4H2O | 1.5 |
| H3BO3 | 3 |
| Na2MoO4 · 2H2O | 2.5 |
| Zn(CH3COO)2 · 2H2O | 13 |
| Fe(II) citrate | 100 |

**Supplementary Table 1. Riesenberg media recipes.** Riesenberg media #1 replicates the batch media recipe listed in DeLisa *et al*. 1999. Trace elements were added together from a single stock solution thus concentrations of all trace element components were increased uniformly. Individual trace element concentrations are listed relative to the entire media.
